# Background food influences rate of encounter and efficacy of rodenticides in wild house mice

**DOI:** 10.1101/2024.01.02.573962

**Authors:** Peter R. Brown, Steve Henry, Lyn A. Hinds, Freya Robinson, Richard P. Duncan, Wendy A. Ruscoe

## Abstract

Baiting is widely used in wildlife management for various purposes, including lethal control, fertility control, disease and parasite control, and conditioned aversion programs for many invasive vertebrate species. The efficacy of baiting programs relies on the likelihood that target animals will encounter the bait, consume it, and receive an appropriate dose of the active ingredient. However, there has been little focus on encounter rate of toxic baits combined with behavioural aversion, which are likely to be significant factors affecting efficacy.
The likelihood of an animal encountering and consuming a toxic grain bait should theoretically increase in proportion to its availability relative to background food quantity if it is neither more or less detectable or palatable. Furthermore, the probability of consuming toxic baits might also be influenced by bait aversion following ingestion of a non-lethal dose of toxin.
Using a model system of wild house mice (*Mus musculus* L.) in mouse-proof enclosures in Australia, we manipulated background food, applied zinc phosphide (ZnP) baits and measured the mortality of mice. When background food was scarce, mouse mortality was high, whereas an increasing abundance of background food led to reduced mortality. A scenario modelling random encounters and including bait aversion explained 78% of the variation in observed mortality outcomes and achieved a closer fit to the data. Mortality rates were predicted to be higher with a higher strength bait which overcomes behavioural aversion.
Ensuring that animals locate and consume a lethal dose of toxic bait is a critical factor for successful bait delivery and efficacy. This is particularly significant in toxic baiting programs, where sublethal doses can make animals feel sick, leading to a negative association with the bait, and the development of aversion.
*Synthesis and applications:* Our findings explain why some toxic baiting programs might fail. To achieve successful control, efforts should be directed at reducing the availability of background food to increase the probability of encounter and uptake of toxic baits. It is important to measure and understand the role of background food on toxic baiting programs to explain variable outcomes and inform strategies for successful bait delivery.

**Graphical abstract:** Our findings explain why some toxic baiting programs might fail. To achieve successful control, efforts should be directed at reducing the availability of background food to increase the probability of encounter and uptake of toxic baits. It is important to measure and understand the role of background food on toxic baiting programs to explain variable outcomes and inform strategies for successful bait delivery.

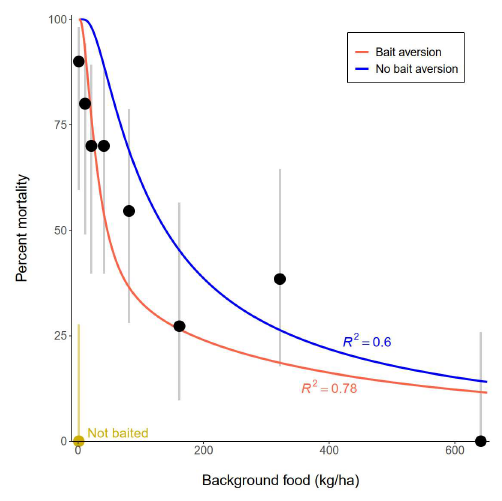

## 1 INTRODUCTION

Baiting is widely used in wildlife management for lethal control and eradications (Aulicky 2022; Capizzi 2020; Marlow *et al*. 2015; Samaniego *et al*. 2021), fertility control (Jacoblinnert *et al*. 2022; Massei and Cowan 2014), disease vaccination (Henning *et al*. 2017; McClure *et al*. 2022), parasite control (Smyser *et al*. 2013), and conditioned aversion programs (O’Donnell *et al*. 2010) of many invasive vertebrate species. The efficacy of baiting programs may be affected by a variety of factors, including bait distribution, attractiveness and palatability, habitat complexity, unintended removal, degradation by environmental conditions (e.g., weather that may affect bait longevity), and the presence of alternative food sources (Allsop *et al*. 2017). Success is determined by the probability that target animals firstly encounter the bait, secondly that they consume the bait, and finally that the bait contains an efficacious dose of the active ingredient (Nugent *et al*. 2011).

During baiting programs, foraging animals are faced with the choice between various food options including the toxic baits. The encounter rate with toxic baits compared with alternative food is important because toxic baits essentially need to compete with other food types for the animal’s attention. To improve the acceptance and/or palatability of baits for lethal control, a toxin is often incorporated into a familiar food item (Buckle and Eason 2015). Toxic baits, such as rodenticides, are often coated on wheat grains (a familiar food type for granivorous rodent pests such as house mice and voles) for use in agricultural settings. The toxic grains are distributed by air or ground onto crops including cereals, canola, lentils, and alfalfa (Brown *et al*. 2002; Jacob *et al*. 2010; Jokić and Blažić 2022) meaning wild mice have to locate the toxic grains in the complex crop environment during foraging activity where non-toxic grains may be abundant. In this situation, we might expect that the probability of an animal encountering and consuming a toxic grain bait will depend on the availability of the similar background food, either maturing seeds/grain on the crop or as grain on the ground due to pre- and post-harvest loss. If we assume that mice do not distinguish between toxic and non-toxic grains and they are equally accessible, then mice will encounter and consume each grain type in direct proportion to their availability. The probability of encountering and consuming a toxic grain (ε) equals the number of toxic grains (*n*_*toxic*_) divided by the total number of toxic and natural/non-toxic (*n*_*non-toxic*_) grains available and declines monotonically as the amount of background non-toxic food increases (Figure 1, solid line).

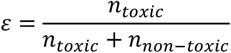

**Figure 1.**
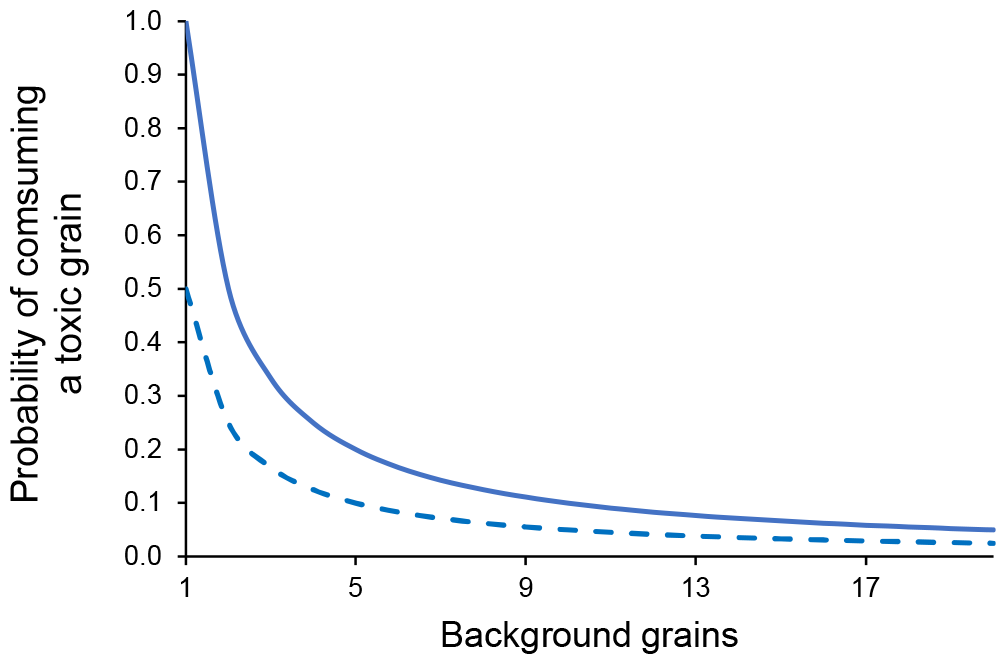
Theoretical probability of an animal finding and consuming one toxic grain amongst an increasing number of background grains given encounter rate is proportional to availability when a) the toxic grain is equally palatable as the non-toxic grain (solid line), and b) the toxic grain is (for example 50%) less acceptable than the non-toxic grain (dashed line).

If the acceptability of the toxic grain is lower than the background food (e.g. lower nutritional quality, general neophobic behaviour of the target animal) we would expect the probability of consumption of a toxic grain to be lower than the encounter rate for any given level of background food as animals decide not to consume the toxic bait in favour of non-toxic food. Bait aversion following a sublethal dose of toxin is one mechanism which is likely to change foraging behaviour when animals associate sublethal toxic side-effects with the bait through appearance, taste, or smell (bait shyness or conditioned aversion) (Allsop *et al*. 2017; Horak *et al*. 2018; Prakash 1988). In this case animals may still encounter baits but their probability of consuming them is much reduced (Figure 1, dashed line) and is related to the level of aversion the sublethal dose has instigated.

The relationship between background food and encounter rate of toxic baits has received little attention but potentially has wide-ranging implications particularly for invasive species management. A notable exception is the use of an understanding of encounter rates to modify bait distribution patterns for the control of invasive brushtail possums in New Zealand (Nugent *et al*. 2012). It is reasonable to assume that when background food is in far greater supply than a toxic bait, that the efficacy will be affected, and although management recommendations often include minimising background/alternative food in the environment (e.g.: Brown *et al*. 2004), very little information exists as to how much of a problem this is, or how little background food can affect toxic bait efficacy.

To explore the importance of encounter rate and sublethally derived bait aversion, we undertook an enclosure trial with wild caught house mice (*Mus musculus*) exposed to the rodenticide, zinc phosphide (Zn_3_P_2,_ hereafter ZnP) as the model system. In Australia, the application of ZnP is a key management tool across large areas of grain-growing regions (Singleton *et al*. 2007) to control invasive house mice but does not always appear to be effective with field studies reporting between 50-95% efficacy (Brown 2006; Brown *et al*. 2002; Mutze and Sinclair 2004; Ruscoe *et al*. 2023b; Twigg *et al*. 2002). This variability could be related to several factors such as the acceptability of the bait substrate (Henry *et al*. 2022; Jacob *et al*. 2010; Johnston *et al*. 2005), mouse behaviour including aversion/bait shyness (Henry *et al*. 2022; Horak *et al*. 2018; Jacob *et al*. 2010; Shepherd and Inglis 1993), the amount of toxin on baits (Hinds *et al*. 2023) and competition with background/alternative food (Mutze 2017). It is often assumed that abundant alternative food will reduce rodenticide efficacy (Brown *et al*. 2002; Guidobono *et al*. 2010), but it is rarely measured (but see Jacob *et al*. 2003). In this system, the alternative or hereafter referred to as background food is mostly grain that has fallen off the standing crop (weather related losses) or has spilled during harvest (harvest losses). In this paper we measure the lethal control of invasive house mice using the registered ZnP bait (25 g ZnP/kg grain) in mouse-specific enclosures where background food quantity is manipulated. This case study examined how expected mortality is a function of three processes:

1. The number of toxic baits that animals (randomly) encounter with varying quantities of background food,
2. The number of baits they consume in a specified period with and without bait aversion occurring, and
3. The mortality rate associated with consuming a given number of toxic baits.

## 2 METHODS

### Enclosure trial

#### 2.1 Mouse enclosures

We set up an enclosure trial to determine the role of background food quantity on mouse mortality from commercially available ZnP25 (zinc phosphide bait; 2.5% ZnP w/w or 25 g ZnP/kg wheat grain) wheat baits. We used enclosures to ensure mouse populations were known (no immigration/emigration), and to control the amount of background food available. We used nine, 225 m^2^ outdoor, predator-proofed mouse enclosures available at the former Mallee Research Station, Walpeup, north-western Victoria, Australia (35°07’S 142°00’E). Enclosures were constructed with galvanised sheet metal buried ∼1 m below ground and to a height of ∼1 m above ground and covered with mesh. The enclosure areas and buffers between fences were mown to remove large weeds and shrubs to achieve a uniform low level of ground cover and to minimise other food sources (for details, see Supplementary Materials).

#### 2.2 Capture of mice for use in enclosures

Mice were captured in a nearby grain farm using Longworth live-capture traps (Longworth Scientific, Abingdon, UK), processed and PIT tagged (10 mm passive integrated transponder, Biomark). Each mouse was randomly assigned to an enclosure, comprising 7 females and 5 males with approximately equal average body weights across all enclosures. Mice were released into enclosures on Day 1 (23 July 2022) for a ten-day acclimation period. We released 12 mice into each enclosure to ensure at least 10 survived the relocation and initial 10 days pre-treatment.

#### 2.3 Application of maintenance food

Sufficient daily maintenance food was provided for each mouse (3 g/day/mouse for Days 1-20) by hand-broadcasting wheat grains randomly but evenly distributed throughout each enclosure (Figure S1, Table 2), such that background food levels remained about constant (see below). Water was provided *ad libitum* at three water stations per enclosure.

**Table 1.** Amount of maintenance food and background food added to each enclosure while the toxic bait was available.

| TREATMENT | | Maintenance food<br>(g per enclosure<br>per day) | Maintenance food<br>(grains per<br>enclosure per day)<br>$n_m$ | Toxic grains<br>(g per<br>enclosure) | Toxic grains<br>(grains per<br>enclosure)<br>$n_p$ |
| --- | --- | --- | --- | --- | --- |
| Added<br>background<br>food (kg/ha) | Added<br>background food<br>(grains per<br>enclosure)<br>$n_a$ | | | | |
| 0 | 0 | 30 | 750 | 22.5 | 562 |
| 10 | 5,625 | 30 | 750 | 22.5 | 562 |
| 20 | 11,250 | 30 | 750 | 22.5 | 562 |
| 40 | 22,500 | 30 | 750 | 22.5 | 562 |
| 80 | 45,000 | 30 | 750 | 22.5 | 562 |
| 160 | 90,000 | 30 | 750 | 22.5 | 562 |
| 320 | 180,000 | 30 | 750 | 22.5 | 562 |
| 640 | 360,000 | 30 | 750 | 22.5 | 562 |
| Control | 0 | 30 | 750 | 0 | 0 |

**Table 2.**
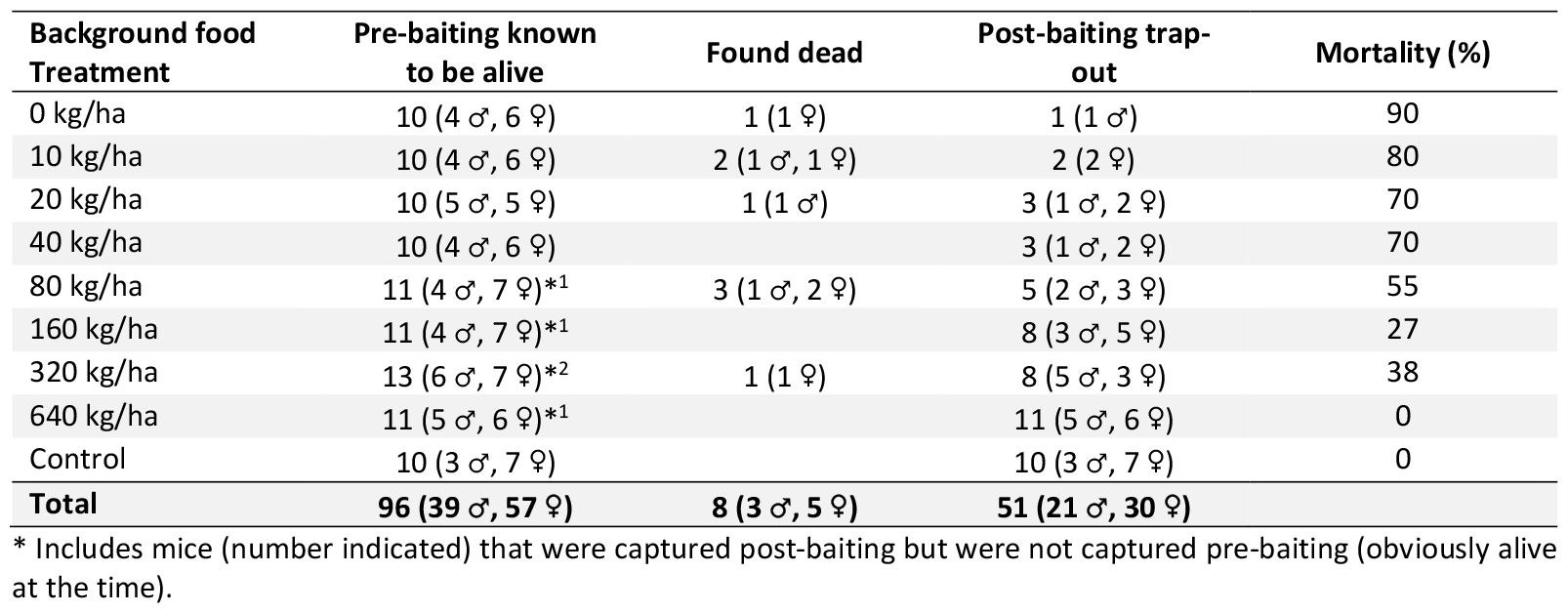
Summary of mouse captures, fate and mortality for each treatment throughout the experiment (♂ = male, ♀ = female). 12 mice were initially introduced into each enclosure but were reduced to a target of 10 mice/enclosure at the pre-baiting trapping. Some mice were subsequently captured during the post-baiting trapping, so were included in the pre-baiting catch. Mortality (%) was calculated by post-baiting catch – pre-baiting catch.

#### 2.4 Application of Treatments: background food

A gradient design (Kreyling *et al*. 2018) was used to detect nonlinear responses to background food availability. On Day 11, background food treatments were applied, and included nil background food (Control enclosure, maintenance food only), then doubling each treatment of added grain from 10 kg/ha through to 640 kg/ha to encompass the likely full range of background food available to mice (Jacob *et al*. 2003) (Ruscoe et al. unpublished data) (Table 2).

#### 2.6 Application of ZnP bait

On Day 16, commercial ZnP25 mouse bait (ZnP25, AG Schilling & Co, Cunliffe, SA) was applied to eight enclosures. Baiting delivered 22.5 g of ZnP coated grain per 225 m^2^ enclosure, equivalent to a rate of 1 kg/ha (approximately 2-3 toxic grains per square metre) as described on the registered label. In the ninth enclosure no ZnP bait was applied (Control). From Day 17-26, each enclosure (n=9) was checked by systematically searching for sick, dying or dead mice under grass clippings or in burrow entrances up to three times a day. Any mouse found was scanned for their PIT tag and weighed. All mice were necropsied to look for signs of ZnP poisoning. Most mice die underground and are not seen.

#### 2.5 Population sampling to assess mortality

Mouse populations were trapped pre- and post-bait application to determine survival (Figure S1). For pre-baiting population sampling, 20 Longworth traps were placed in each enclosure for three nights (Days 13-16; total = 60 trap nights/enclosure; Figure S1). The number of mice per enclosure was reduced in some enclosures but not to less than 10 (∼450 mice per ha) during the pre-baiting population sampling (not all animals were recaptured during this period). This number reflects the moderate mouse densities at which growers are trying to control mice to minimise crop damage and was considered the minimum needed to adequately observe treatment effects and provide statistically robust data.

Trapping effort was increased for the post-baiting population sampling (33 Longworth traps per enclosure, removal trapping; Figure S1). Trapping ran for six nights (Days 20-26; ∼200 trap nights per enclosure), and on the last two nights, cornflour was dusted over likely mouse burrows to look for active burrows where additional traps were laid. Trapping continued until no mouse activity was observed for two days (as assessed by no captures and undisturbed cornflour on burrows). Captured mice were scanned for PIT tags, weighed, head-body length measured and examined for overall body condition and signs of poisoning. Mice were then humanely killed by cervical dislocation and necropsied to assess the condition of major organs (liver, spleen, kidneys, stomach content, lungs) to record sub-acute signs of toxicity.

For analysis, we included all animals that were trapped during the pre- and post-baiting trapping only (where we had a known fate; when numbers were reduced to ten individuals per enclosure). We were unable to determine the fate of a small number of mice (n = 9), because they were not re-captured at any time after they were introduced into the enclosures on Day 0 (when 12 mice were introduced into the enclosures), so we have assumed they died underground during the acclimation period.

#### 2.7 Bait toxicity testing

To verify bait toxicity, a sample of ZnP25 grains was randomly taken from the same drum used to bait the enclosures and individually analysed (n=20 grains) by ACS Laboratories [Australia] Pty Ltd (Kensington, Victoria, Australia) (see Supplementary Material for details).

### Predicted mortality

We used independent data on mouse survival as a function of the number of toxic grains eaten (Section 2.8), coupled with modelled bait encounter rate estimates to predict mouse mortality in relation to background food availability (Section 2.9), and then compared these predictions to the outcomes observed in the enclosure trial. We predicted outcomes under two scenarios: (1) mice randomly encountered and consumed toxic grains with no bait aversion (Section 2.10), and (2) mice randomly encountered toxic grains but became averse to consuming toxic grains if they had previously consumed them but not consumed a lethal dose (Section 2.11).

#### 2.8 Survival probability based on number of toxic grains eaten

We obtained data on mouse survival as a function of the number of toxic grains eaten from two laboratory studies: Henry *et al*. (2022) and Hinds *et al*. (2023). The Henry *et al*. (2022) study reported data on mouse survival following consumption of ZnP25 grains (n = 90 mice) (25 g ZnP/kg grain bait), while the Hinds *et al*. (2023) study reported data on mouse survival following consumption of both ZnP25 grains (n = 13) and ZnP50 grains available under a temporary use permit (PER90799) (50 g ZnP/kg wheat grain bait, n = 24). For ZnP25, we combined data from both studies to estimate survival rate.

For a given total number of toxic grains eaten *T*_p_, we had data on the number of mice *N* that had consumed that number of grains, and the number of mice alive *S* at the end of each trial. We fitted separate exponential survival models to the ZnP25 and ZnP50 data of the form:

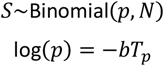

Where *p* is the probability of survival, *b* is the exponential survival parameter – which took a different value for ZnP25 and ZnP50 grains – and log is the natural logarithm. We fitted the models to the data in a Bayesian framework using JAGS v4.3.1 implemented through the package jagsUI in R v4.2.1. Note that there was variation in the numbers of mice that consumed different numbers of grains in the laboratory trials. The binomial model we fitted accounts for this variation in sample size (see Figure 2).

**Figure 2.**
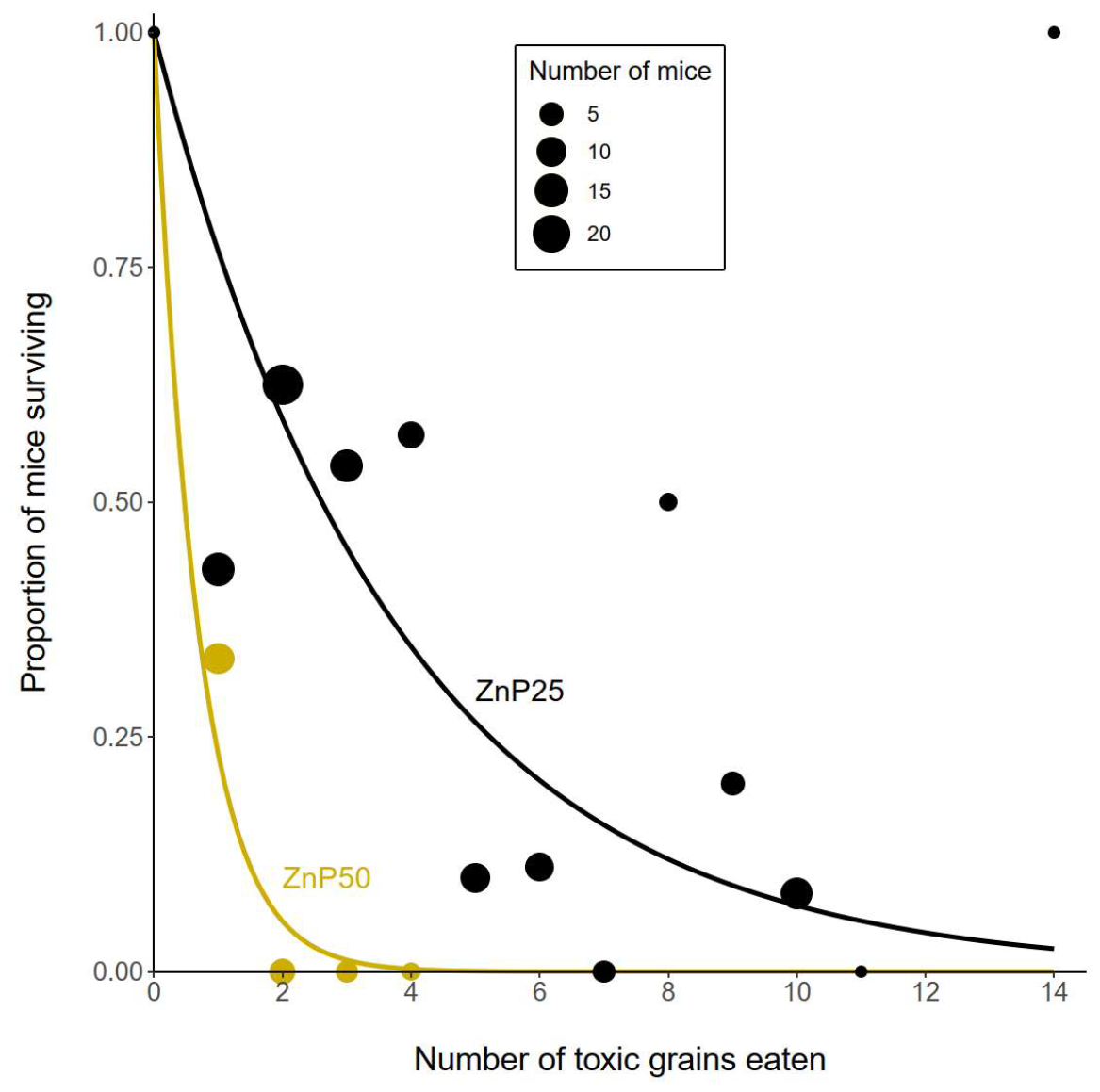
Proportion of mice surviving as a function of the number of toxic grains consumed for ZnP25 (black filled circles) and ZnP50 (gold filled circles) grains. Circle size is proportional to the number of mice used to estimate the survival proportions. Solid lines are the exponential survival curves fitted to the data. Data sourced from Henry *et al*. (2022) and Hinds *et al*. (2023).

#### 2.9 Bait encounter probability

Maintenance food was replenished each day, but toxic grains and non-toxic background food were not. Therefore, we expect bait encounter probabilities to change over time as grains are consumed within each enclosure. Table 1 shows the starting grain availability in each enclosure at the time toxic grains were added, calculated using an average grain weight of 0.04 g (based on measuring the weight of 45 grains, standard deviation = 0.011) and enclosure area of 0.0225 ha.

#### 2.10 Encounter probability with no bait aversion

In this scenario we assume that mice do not distinguish between toxic and non-toxic grains, such that they encounter and consume each grain type in direct proportion to their availability. At the start of night 1 (Day 16), following addition of toxic grains to each enclosure, the total number of non-toxic grains *n*_*b*,1_ is the sum of the added background food and the daily maintenance food *n*_*b*,1_ = *n*_*a*_ + *n*_*m*_. The number of toxic grains *n*_*p*,1_ = *n*_*p*_ where values for *n*_*p*_, *n*_*a*_ and *n*_*m*_ are given in Table 1. On night 1, the probability that a randomly chosen grain is a toxic grain is then:

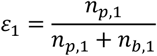

If there are *M*_1_ mice per enclosure on night 1 and each mouse consumes *c* grains per night, then total grain consumption on night 1 is *M*_1_*c*. The expected number of toxic grains consumed per mouse on night 1 *c*_*p*,1_ = *ε*_1_*c*, and the total number of toxic grains consumed that night is *M*_1_*c*_*p*,1_. The expected number of non-toxic grains consumed per mouse on night 1 *c*_*n*,1_ = (1 − *ε*_1_)*c*.

As a consequence of consuming toxic grains on night 1, some mice could die and hence would not consume any grains on night 2 (Day 17). If each mouse was expected to consume *c*_*p*,1_ toxic grains, then, from the survival curve, the number of mice surviving to night 2 is: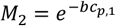.

At the start of night 2, the expected number of toxic and non-toxic grains in an enclosure given consumption on night 1 and the addition of maintenance food is:

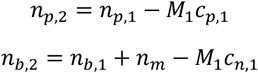

And the probability that a randomly chosen grain is a toxic grain is:

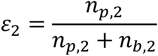

The expected number of toxic grains consumed per mouse on night 2 *c*_*p*,2_ = *ε*_2_*c*, and the number of mice surviving to night 3 is: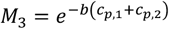.

Extending this to any night *i* > 1, the number of toxic and non-toxic grains present at the start of night *i* following consumption during night *i* − 1 and the addition of maintenance food is:

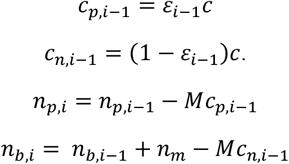

So that:

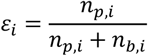

And

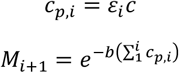

We can obtain toxic grain encounter probabilities for consecutive nights by iterating these equations and can calculate from these the expected number of toxic grains consumed per mouse each successive night *c*_*p,i*_ and the resulting number of surviving mice *M*_*i*_. We can then calculate the expected mortality rate *D* for a given level of background food following *n* nights of exposure as:

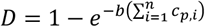

#### 2.11 Encounter probability with bait aversion

In this scenario we assume that if a mouse consumes a toxic bait and survives, it is less likely to consume toxic baits in the future. We model this aversion by assuming that having consumed a toxic bait, only some proportion *λ* of mice will continue to consume toxic baits by making no distinction between these and non-toxic baits. A proportion 1 − *λ* of mice that consume a toxic bait will be averse to those baits and stop consuming them.

We assume that bait aversion occurs the night after a mouse consumed one or more toxic grains, given it takes time for an adverse response to occur. Consequently, for night 1 the encounter probability and number of toxic grains consumed will be the same as for the no bait aversion scenario above. However, some mice are expected to have consumed toxic grains during night 1 and are thus less likely to consume toxic grains during night 2. The probability a mouse did not consume a toxic grain during night 1 is (1 − *ε*_1_)^*c*^ and the probability it did consume a toxic grain is (1 − (1 − *ε*_1_)^*c*^).

The expected number of toxic grains consumed by a mouse during night 2 is the probability of encountering a bait on night 2, *ε*_2_*c*, multiplied by the probability of consuming that bait, which equals the sum of the probability that a mouse did not consume a bait on night 1 and the probability that a mouse did consume a bait on night 1 multiplied by *λ*: *c*_*p*,2_ = *ε*_2_*c*[(1 − *ε*_1_)^c^ + *λ*(1 − (1 − *ε*_1_)^*c*^)]. The expected number of non-toxic grains consumed by a mouse is then *c*_*n*,2_ = *ε*_2_*c*(1 − [(1 − *ε*_1_)^*c*^ + *λ*(1 − (1 − *ε*_1_)^*c*^)]). For night 3, the expected number of toxic grains consumed is the probability of encountering a bait on night 3 multiplied by the probability of consuming that bait, which equals the sum of the probability that a mouse did not consume a bait on nights 1 or 2 plus the probability that a mouse did consume a bait on nights 1 or 2 multiplied by *λ*: *c*_*p*,3_ = *ε*_3_*c*[(1 − *ε*_1_)^*c*^(1 − *ε*_2_)^*c*^ + *λ*(1 − (1 − *ε*_1_)^*c*^)(1 − (1 − *ε*_2_)^*c*^)], and so forth for subsequent nights. For the bait aversion scenario, we can thus calculate the expected number of toxic grains consumed by a mouse each night, calculate the number of mice surviving after each night, and obtain the expected mortality after *n* nights as for the no bait aversion scenario above.

#### 2.12 Parameter estimates

To calculate expected mortality given initial background food availability *n*_*a*_ + *n*_*m*_ and number of toxic grains *n*_*p*_ using the above approach requires specifying values for 5 parameters: the exponential survival parameter *b*, calculated for ZnP25 toxic grains using the data and methods described above; the number of mice per enclosure *M*∼10; the number of nights during which mice were exposed to toxic baits *n* = 5, the number of nights between the addition of toxic grains and post-baiting trapping; the number of grains consumed per mouse per night *c* = 75 (calculated assuming each mouse consumes 3 g of food per night with an average grain weight of 0.04 g); and the proportion of mice that did not become averse to toxic grains following consumption *λ* = 0.2, obtained from laboratory feeding trial data in Henry *et al*. (2022).

Using these parameter estimates, we calculated expected mortality given a range of background food availability in enclosures ranging from 0 to 640 kg/ha for the no aversion and bait aversion scenarios using the methods described above. We compared the mortality predicted under each scenario with mortality observed in the field trial by calculating the residual sum of squares *RSS* = ∑(*observed* − *predicted*)^2^ and used this to calculate an *R*^2^ value for each scenario specifying the proportion of variation in observed mortality rates explained by the predictive model: 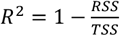, where *TSS* is the total sum of squares of the observed mortality rates.

The field trial used ZnP25 toxic grains, so we predicted mortality using the survival parameter *b* derived from laboratory trials that used ZnP25 toxic grains. In addition, we calculated predicted mortality using the survival parameter derived from laboratory trials that used ZnP50 toxic grains (Hinds *et al*. 2023) to predict the relationship between background food availability and mouse mortality using ZnP50 baits.

## 3 RESULTS

### 3.1 Enclosure mortality

Mouse mortality ranged from 0% (640 kg/ha and Control enclosures) to 90% (0 kg/ha) (Table 2). Mortality was high when there were low levels of background food present (mortality ≥70% for 0-40 kg/ha background food), but mortality was low when there were abundant levels of background food present (mortality <40% for >160 kg/ha background food). In order to achieve >70% mortality, background food needs to be less than 40-80 kg/ha.

### 3.2 Survival probability based on number of toxic grains eaten

Figure 2 shows the laboratory data of Henry *et al*. (2022) and Hinds *et al*. (2023) used to model survival probability as a function of the number of toxic grains consumed, and model fits of the exponential survival curves for ZnP25 and ZnP50 grains. For ZnP25 grains the exponential survival parameter *b* = 0.265 with 95% credible intervals 0.199-0.341. For ZnP50 grains the exponential survival parameter *b* = 1.474 with 95% credible intervals 0.803-2.423.

### 3.3 Expected number of toxic baits consumed

Figure 3 shows the expected number of toxic grains consumed per mouse over five nights as a function of background food availability for the two modelled scenarios: with no bait aversion and with bait aversion. In both scenarios, the number of toxic grains consumed declines markedly as background food availability increases above zero, noting that the y axis of Figure 3 is a log scale.

**Figure 3.**
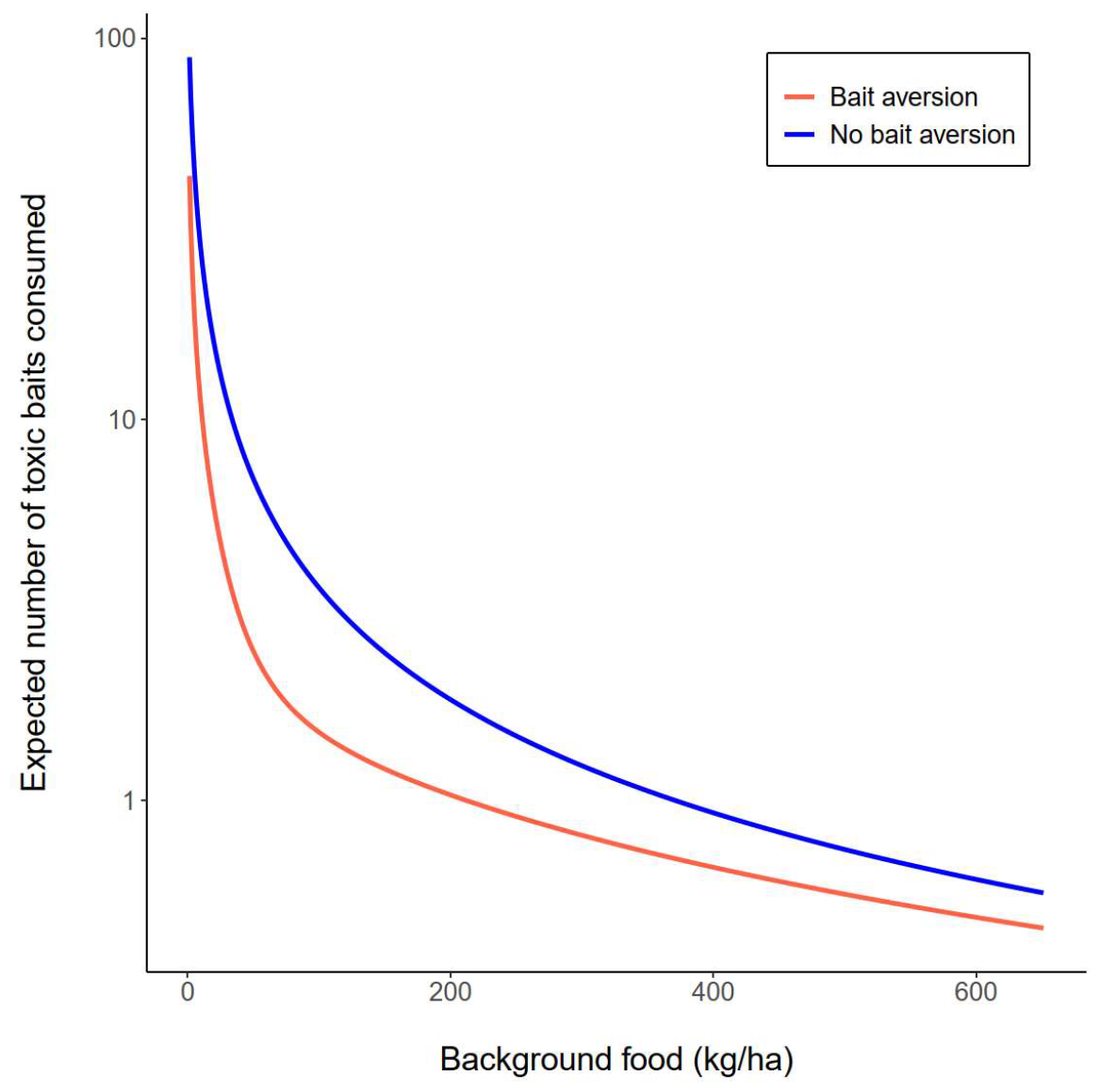
Expected number of toxic grains consumed per mouse over five nights (on log scale) as a function of background food availability for the two modelled scenarios: with no bait aversion and with bait aversion.

### 3.4 Observed and predicted mortality rates

There was zero mouse mortality in the Control enclosure (maintenance food only provided; no toxic bait added) (Figure 4). In the enclosures with toxic grains added, mouse mortality rates were highest (>75%) in enclosures with the lowest background food availability, and mortality rate generally declined as background food availability increased.

**Figure 4.**
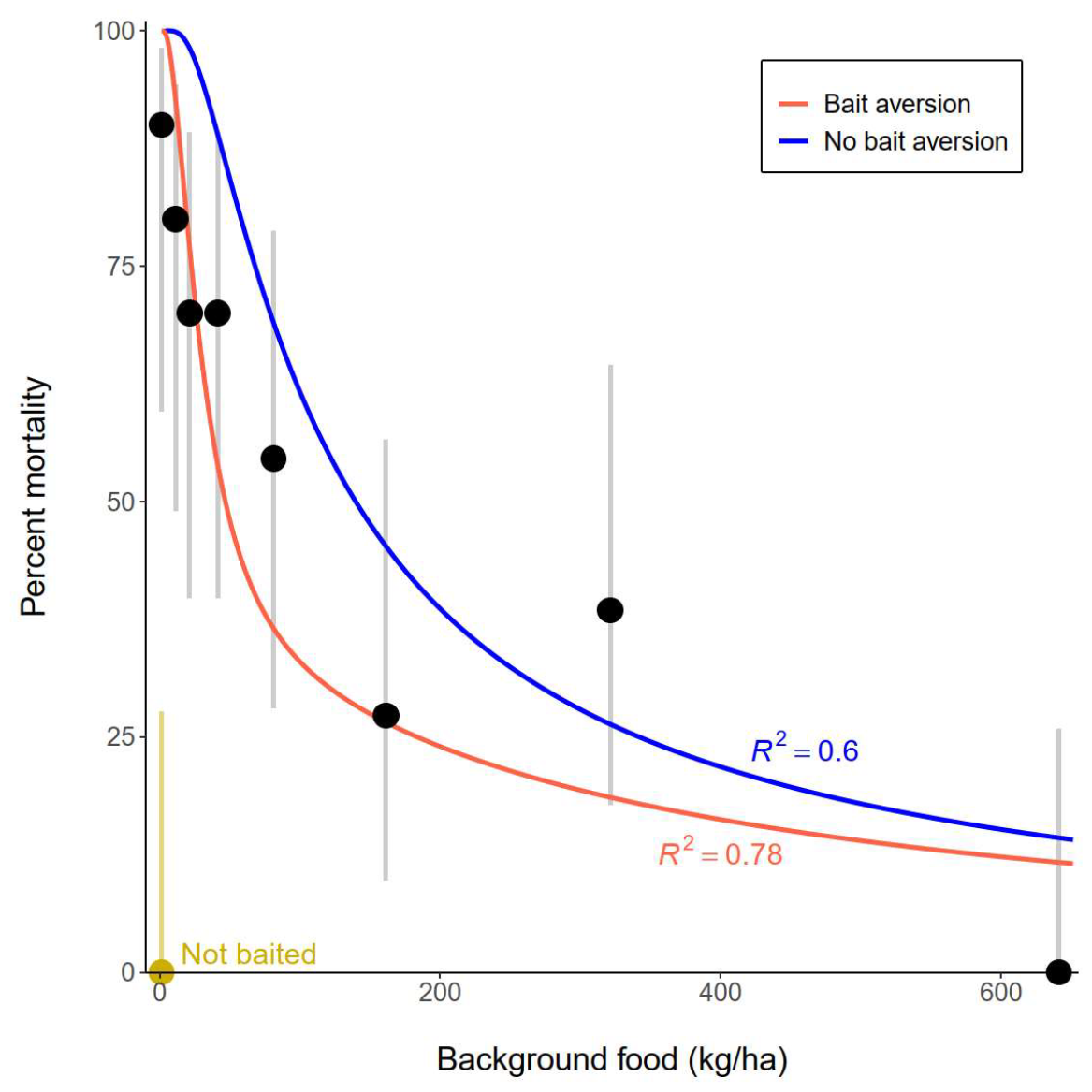
Observed mouse mortality as a function of background food availability for enclosures with added toxic grains (black filled circles) and the control enclosure with no toxic grains (gold filled circle) and associated 95% confidence intervals (grey and gold lines, calculated for binomial data using the Wilson method). Solid lines show the mortality rates predicted under the two modelled scenarios: random bait encounters without bait aversion, and random bait encounters with bait aversion.

Consistent with the decline in the expected number of toxic grains consumed (Figure 3), predicted mortality declined with increasing food availability in both the no bait aversion and bait aversion scenarios (Figure 4). The scenario modelling random encounters with no bait aversion could explain 60% of the variation in observed mortality outcomes, but tended to consistently overestimate rates of mortality, particularly at low levels of background food availability where predicted outcomes fell outside the 95% confidence intervals of the observed outcomes. In contrast, the scenario modelling random encounters with bait aversion provided a better fit to the data, explaining 78% of the variation in observed mortality outcomes and achieving a closer fit to the data at low levels of background food availability; predicted outcomes were inside the 95% confidence intervals of the observed outcomes for all but the zero level of background food availability.

### 3.5 Predicted outcomes for ZnP50 baits

The laboratory trials show that, for a given number of grains eaten, ZnP50 baits cause higher mortality (lower survival) than ZnP25 baits (Figure 2). Consequently, for a given level of background food availability, mortality rates are predicted to be higher for ZnP50 relative to ZnP25 baits for both scenarios (Figure 5). While mortality rates are also predicted to decline with increasing background food availability for ZnP50 baits, this decline is less marked than for ZnP25 baits. At the highest level of background food availability (640 kg/ha), mouse mortality with ZnP50 grains and bait aversion is predicted to be >50%, compared to just over 10% for ZnP25 baits (Figure 5).

**Figure 5.**
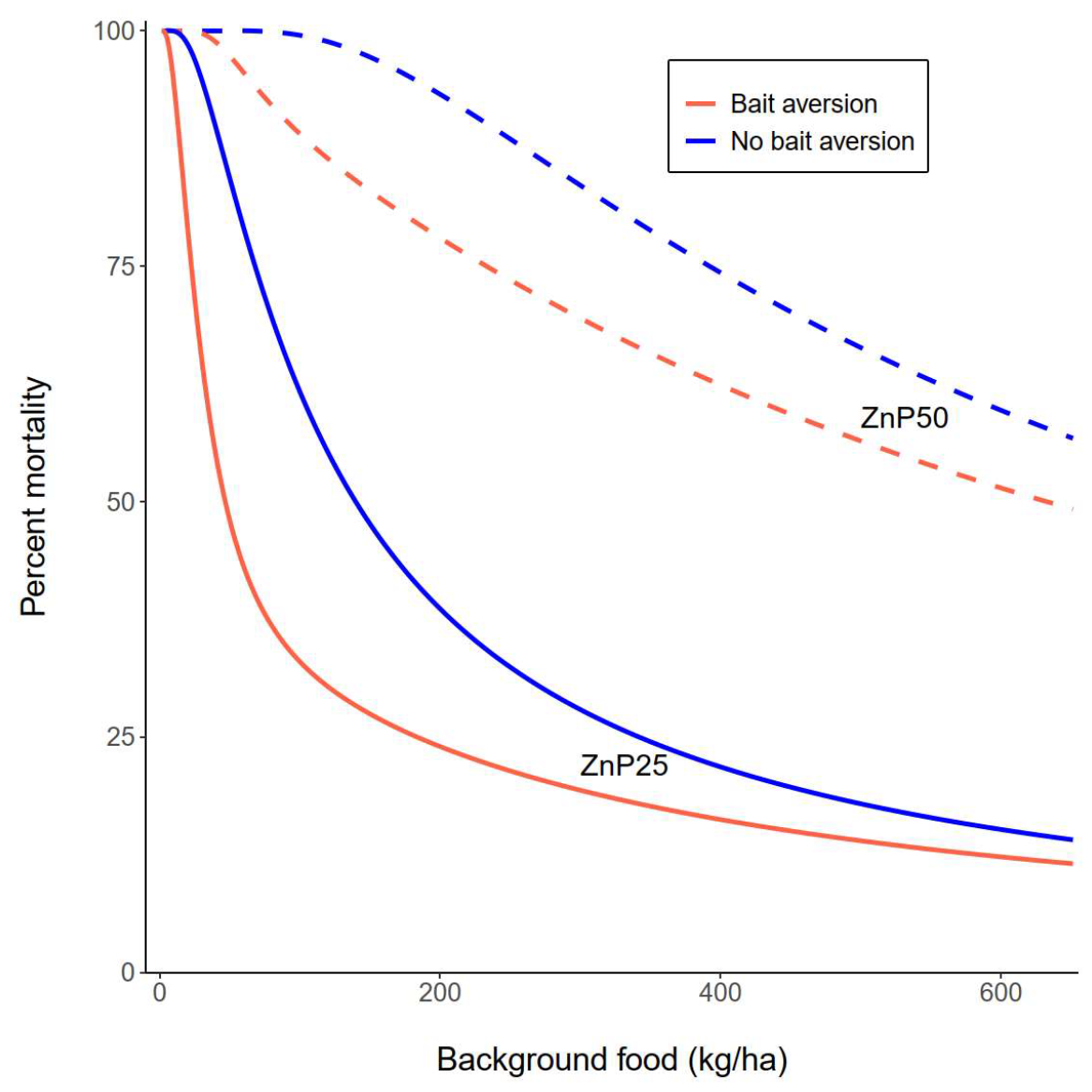
Mortality rates predicted under the two modelled scenarios: random bait encounters without bait aversion, and random bait encounters with bait aversion with either the ZnP25 bait as used in the field trial, or with the newer ZnP50 bait.

### 3.6 Necropsy of mice found dead

A total of eight mice were found in all enclosures over the 11 days of checking (Table 1); which is 17.8% of mice assumed dead (8/45 dead mice). All found mice were detected when conducting visual assessments during the day. No dead mice were found in the Untreated Control enclosure, or enclosures for 40 kg/ha, 160 kg/ha and 640 kg/ha.

On necropsy, all eight mice found dead showed signs of mottling and colour change in their livers, consistent with ZnP-affected mouse livers observed in lab studies and were considered to have died from zinc phosphide poisoning (Table S2). Their kidneys, spleen, heart and lungs were all normal size and colour. There were 51 mice re-captured in the post-baiting population sampling and removed, measured and necropsied for signs of ZnP poisoning. There were no overt signs of ZnP poisoning in these mice (all organs were considered “normal”).

### 3.7 Bait testing

The toxicity of the bait used in the trial was 1.05 mg ZnP/grain (± 0.28 SD, CV = 5.55%, n=20; range 0.7-1.9 mg/grain), which is equivalent to 25 g/kg bait (ZnP25).

## 4 DISCUSSION

Two critical factors that impact the efficiency of toxic baits are encounter rate (which is influenced by bait distribution and availability of background food) and bait aversion as a result of consuming a sublethal dose. While these factors are generally assumed, we have empirically demonstrated the effect of these for house mice in enclosures. Our findings demonstrated that when abundant wheat grain was present, mice were less likely to encounter and consume rodenticide-coated wheat grain bait and die. Even at relatively low levels of this background food, the effectiveness of ZnP baiting decreased rapidly. These findings have significant implications for the management of house mice in Australian cropping systems, as well as for pest management and conservation efforts more generally. These factors are critical when designing control or management strategies for other types of baits, including contraceptives, and are not limited to just rodenticides. These findings might help to explain why other control programs may not have been successful (Guidobono *et al*. 2010; Jacob *et al*. 2003; Samaniego *et al*. 2022; Samaniego *et al*. 2023).

In the context of using rodenticides against mice, the challenge lies in the probability of mice discovering the toxic baits in the presence of background food. Mice in this enclosure study chose between wheat grains (background food) and ZnP-coated toxic wheat grains. It is likely that individual mice exhibit different foraging behaviours, influenced by factors such as seed dispersion patterns, seed size, handling time, moisture, energy and soluble carbohydrate content of the seeds, and secondary chemical compounds like polyphenols (Kerley and Erasmus 1991). One question that arises is whether the rodenticide baits need to be more attractive to increase encounter rate, or if mice simply sample all food they encounter, as we have assumed here. This raises considerations regarding the encounter rate of the baits and the principles of optimal foraging theory. Optimal foraging theory suggests that mice should select a subset of food from a set of potential food to maximise net energy intake per unit time spent foraging (Charnov 1976; Pulliam 1974). Our results suggest that when abundant similar food sources are available, mice do not invest significant time foraging for food, which reduces their likelihood of encountering a toxic grain amongst higher densities of non-toxic grain. When limited background food is available, the chance of encountering and consuming toxic grain is much higher compared to situations with abundant background food.

A critical factor in a baiting program is ensuring that animals encounter and consume a bait in a timely manner. This is particularly significant in toxic baiting programs, as consuming a sublethal dose can make animals feel sick, leading to a negative association with the bait, and the development of aversion (Horak *et al*. 2018; Prakash 1988). To prevent toxin-aversion, either every bait must contain a lethal dose, or baits must be strategically placed to ensure multiple baits are encountered and consumed before sublethal effects of the toxin are experienced by the target animal. Research on baiting of the invasive possum, *Trichosurus vulpecula*, in New Zealand has demonstrated that distributing baits in clusters or strips, rather than uniformly, can increase encounter rate, therefore efficacy, and reduce the total amount of toxin required (Nugent *et al*. 2012). When levels of background food are low, random encounter rates may be sufficient to achieve high baiting efficacy. However, as the amount of background food increases animals become satiated and stop foraging before an adequate number of toxic baits are encountered and consumed. Ensuring every single toxic bait consumed is lethal appears to be important, as we have revealed in our modelling.

Recent laboratory and field studies have demonstrated increased efficacy of the ZnP50 bait over the currently registered ZnP25 bait (Hinds *et al*. 2023; Ruscoe *et al*. 2023b). Our modelling, which considered encounter rate, bait consumption (while accounting for behavioural aversion) and mortality rate, further corroborates this. Moreover, our models indicate that the ZnP50 bait offers increased efficacy in the presence of higher levels of background food because it overcomes bait aversion issues (each bait is a lethal dose). For instance, when background food reaches 300 kg/ha, the ZnP50 bait may still achieve ∼70% mortality, surpassing the ZnP25 bait’s mortality of ∼20% (Figure 5). A gap in our knowledge is the duration of aversion, which will influence follow-up management.

The issue of competing food resources and/or encounter rate are important in other situations. For example, the toxic bait for invasive feral cat, *Felis catus*, in Australia is best applied into resource-poor environments when lower densities of prey items (native small mammals, birds and reptiles) mean the cats have to travel greater distances to forage (Tiller *et al*. 2021). In such cases, toxin-laced meat baits were more likely to be encountered as they were non-mobile and can be placed at likely areas of activity (water holes, tracks) of the pest predator. Doucette (1954) found that bait placement for black slugs (*Arion ater*, Linnaeus) was critical with snail bait attractiveness (and effectiveness) reduced in the presence of plant foliage compared to fallow ground. Mortality was increased by increasing encounter rates by placing the toxin along the slug’s nocturnal migration routes.

Rodents are a worldwide problem in plant production (agriculture, forestry), public health (intensive livestock production, urban environments, food processing facilities), infrastructure, and conservation (Buckle and Smith 2015; Diagne *et al*. 2023; Howald *et al*. 2007; Meerburg *et al*. 2009; Singleton *et al*. 2021; Swanepoel *et al*. 2017). It is crucial to implement control measures that are highly effective while causing minimal harm to the environment.

In Australia, most mouse damage to winter crops occurs at sowing which coincides with the end of the mouse breeding season when numbers are high (Brown *et al*. 2007; Brown and Singleton 2002), but background food should be relatively low (because it is several months since the previous crop harvest). Jacob *et al*. (2003) found that spilled grain after harvest declined over time from a peak of 206 kg/ha immediately after harvest (December), to 146 kg/ha in January and to 36 kg/ha in February. This decline occurs as food is consumed by mice and other animals and as grain germinates during summer rains. However, our work has shown that even at a relatively low level of background food the effectiveness of the bait declines. Given ZnP is broadcast on the ground at 1 kg/ha (equivalent of 2-3 grains/m^2^), it is unlikely that mice will encounter ZnP-treated grain in amongst existing spilt grain during nocturnal foraging. Even if application rates were to exceed 1 kg/ha (as has been allowed under emergency permits) the dilution against abundant background food availability is still significant. Provided there is “negligible” grain left on the ground when planting new crops, there is a high likelihood that the application of an effective rodenticide should be successful. In conservation farming systems, where grain losses (pre- and post-harvest) are often high, the challenge is to define strategies to reduce either the amount of background food or the availability of the background food prior to application of bait. Additional farm management practices to reduce background food accessibility post-harvest include (based on Brown *et al*. 2004) (1) reducing spilled grain losses during harvest, (2) using seed destructors to remove weed seeds (Walsh *et al*. 2012), (3) using livestock (eg: sheep) to graze post-harvest stubble and grain, and (4) treating post-harvest stubble in some way (burn, prickle-chain etc) (Ruscoe *et al*. 2023a). Further research is needed to determine the benefits of these practices in reducing mouse food supply.

To achieve a 70% mortality in our ZnP enclosure trial, background food needed to be less than about 80 kg/ha (or ∼30 kg/ha assuming aversion was occurring, Figure 4). Although the response curve was an asymptotic decay-shaped curve, it was generally linear through the range of 20-160 kg/ha: incremental increases in background food corresponded to reduced mortality. Modelling the use of ZnP50 improved baiting effectiveness compared to the ZnP25 bait as each toxic grain was lethal and aversion was minimised. Using ZnP50, we estimated that 70% mortality would be achieved when background food was as high as 450 kg/ha (or ∼300 kg/ha incorporating aversion, Figure 5).

## 5 CONCLUSIONS

Our study highlights the critical role of background food in the efficacy of toxic bait use and emphasises the importance of ensuring each bait is a lethal dose to minimise sublethal dosing and subsequent bait aversion. This finding has significant implications for mouse management in Australia, most importantly that rodenticide baits should be applied when available background food is minimal. Further investigations should be conducted in different farming systems, and with other rodent pests to determine if similar patterns exist elsewhere. While toxic baiting can be very effective at controlling pests, additional ecologically-based rodent management strategies (Capizzi *et al*. 2014; Singleton 1997; Singleton *et al*. 2007; Singleton *et al*. 1999), including practices that target the reduction of background food sources are warranted.

## Author contributions

PRB, WAR, LAH and SH designed this research. All authors collected experimental field data. PRB, RPD and WAR analysed the data. PRB, RPD and WAR wrote the first draft manuscript. All authors contributed to the final manuscript.

## Acknowledgements

We sincerely thank the farmers for allowing us to collect mice for this experiment, and to the Victorian Department of Environment, Land, Water and Planning for approval to conduct the experiment at the enclosures. We thank Michael Davies for assistance prior to experimental field work and Peter Jones for assistance in preparation of the enclosures. We thank Leigh Nelson and Ken Young (GRDC) for ongoing support.

## Ethics approval

This study was approved by the CSIRO Wildlife and Large Animal, Animal Ethics Committee (approval number: 2021-30) and adheres to the 8th Edition of the Australian Code and Use of Animals for Scientific Purposes. This article does not contain any studies with human participants performed by any of the authors.

## Data availability

Data are held by the Commonwealth Science and Industry Research Organisation (CSIRO), Australia, and will be made available upon reasonable request to the corresponding author.

