## Supplementary file for "Background food influences rate of encounter and efficacy of rodenticides in wild house mice"

**Supplementary Materials**

**Detailed field methods**

1. Field trial

*Mouse enclosures*

We set up an enclosure trial to determine the role of background food quantity on mice mortality from commercially available ZnP wheat baits. We used enclosures to ensure mouse populations were closed (no immigration/emigration) and to control the amount of background food available. This enclosure experiment was set up in nine outdoor, predator-proofed mouse enclosures at the former Mallee Research Station, Walpeup, in the central mallee wheatlands, north-western Victoria (35°07'26.4"S 141°59'59.0"E). The construction of these enclosures has been described in Barker et al. (1991) with galvanised sheet metal buried ~1 m below ground and to a height of ~1 m above ground. Each enclosure measured 15 x 15 m (225 m2), with a buffer zone (2 m wide) to a second mouse-proof fence. The enclosures were predator-proofed using galvanised wire netting (5 cm mesh, 1 mm wire) constructed to a height of 2.5 m around each enclosure. The netting also excluded aerial predators and grain-eating birds (pigeons, parrots etc.). The enclosure areas and buffers between fences were mown or slashed to remove large weeds and shrubs to achieve a uniform low level of ground cover and to minimise other food sources.

*Capture of mice for use in enclosures*

Longworth live-capture traps (Longworth Scientific, Abingdon, UK) were set in a nearby grain farm (~3 km away) to capture the required number of mice required (108; 12 mice for 9 enclosures). Captured mice were taken back to a central processing area for initial processing. Each mouse was PIT tagged (10 mm passive integrated transponder, Biomark), sexed, measured for length and weight. Each mouse was randomly assigned to an enclosure, and checked to ensure a balance of 7 females and 5 males with approximately equal average body weights across enclosures. Mice were released into enclosures on Day 1 (23 July 2022) for a ten-day acclimation period. We released 12 mice into each enclosure initially to ensure at least 10 survived the relocation and initial 10 days pre-treatment.

*Application of maintenance food*

Daily maintenance food was provided to each enclosure by hand-broadcasting wheat grains randomly and evenly throughout each enclosure (36 g/day for Days 1-16 for 12 mice per enclosure, then 30 g/day for

Days 17-20 for 10 mice per enclosure, Figure S1, Table 2). Water was provided *ad libitum* at three water stations (1.5 L Pet One poultry gravity drinker) at regular spacing throughout each enclosure.

| Day | -4 | -3 | -2 | -1 | 0 | 1 | 2 | 3 | 4 | 5 | 6 | 7 | 8 | 9 | 10 | 11 | 12 | 13 | 14 | 15 | 16 | 17 | 18 | 19 | 20 | 21 | 22 | 23 | 24 | 25 | 26 |
|---|---|---|---|---|---|---|---|---|---|---|---|---|---|---|---|---|---|---|---|---|---|---|---|---|---|---|---|---|---|---|---|
| Trap mice | | | | | | | | | | | | | | | | | | | | | | | | | | | | | | | |
| Maintenance diet (added daily) | | | | | | | | | | | | | | | | | | | | | | | | | | | | | | | |
| Acclimation period | | | | | | | | | | | | | | | | | | | | | | | | | | | | | | | |
| Added background food present | | | | | | | | | | | | | | | | | | | | | | | | | | | | | | | |
| Pre-Treatment population sampling (n = 20 traps/night) | | | | | | | | | | | | | | | | | | Set | | | PU | | | | | | | | | | |
| Zinc phosphide bait applied (at 1 kg/ha) | | | | | | | | | | | | | | | | | | | | | | | | | | | | | | | |
| Monitoring for sick or dead mice (n = number of checks/day) | | | | | | | | | | | | | | | | | | | | | | 3 | 2 | 3 | 2 | 2 | 2 | 2 | 2 | 2 | 1 |
| Post-Treatment population sampling (n = 33 traps per night) | | | | | | | | | | | | | | | | | | | | | | | | | Set | | | | | | PU |

**Figure S1**. Schedule of activities undertaken for enclosure trial to test effect of background food on mortality of mice after zinc phosphide baiting. Mice were sourced from nearby grain farms and 12 mice (7 females, 5 males) were introduced to enclosures on Day 1. Maintenance food (36 g per enclosure) was provided each from Day 1 to Day 20. Background food (Treatment) was provided once on Day 11 (see Table 1 for details of amount provided to each enclosure). Pre-baiting population sampling was conducted Days 13-16, where the number of mice was reduced to 10 mice (6 females and 4 males), baiting occurred on Day 16 (at 1 kg/ha ZnP) and post-baiting population sampling (and removal) was conducted Days 20-26 ("Set" = traps opened, "PU" = traps picked up). Inspections were conducted in each enclosure starting in the day after bait was applied at various times of the day to look for sick and dying mice.

*Application of ZnP bait*

Commercial ZnP mouse bait was used (AG Schilling & Co, (Cunliffe, SA), Mouse bait – sterilised – 25 g/kg, batch#CGF/2022/B1946SX; ZnP25). For each enclosure receiving bait (all except the Control enclosure), 22.5 g of ZnP bait was weighed out in plastic bags (equivalent to a rate of 1 kg/ha as described on the label rate but applied to the enclosure area of 225 m2). Bait was distributed on Day 16 (Figure S1) into small plastic cups to facilitate handling by staff and was applied by hand wearing full PPE (gloves, masks). Quadrats of 1 $m^2$ (1 x 1 m) were used to assist with regular placement of grains of ZnP bait. Working in a team of 2 or 3 staff, the quadrats were placed on the edge of the enclosure and either 2 or 3 grains were randomly placed in each quadrat. The quadrat was then flipped over to the next 1 m interval, and either 2 or 3 grains were randomly placed in each quadrat. This was repeated until the entire area of the enclosure was covered. Any remaining grains of bait were then randomly spread across of the enclosure such that 22.5 g of bait were applied as evenly as possible throughout each enclosure.

Small amounts of rainfall occurred most days after bait application (Table S1), but considered small enough to not affect the toxicity of the ZnP baits.

**Table S1**. Daily rainfall (mm) measured at Walpeup (BOM Station 076064). ZnP baits were applied on Day 16. Days 0 -10 were the initial acclimation period.

| Day number | Date | Activity | Rainfall |
|---|---|---|---|
| Day 11 | 2 Aug 2022 | Background (Treatment) food added | 0 mm |
| Day 12 | 3 Aug 2022 | | 1.6 mm |
| Day 13 | 4 Aug 2022 | Traps set (Pre-baiting population assessment) | 0 mm |
| Day 14 | 5 Aug 2022 | Check traps day 1 | 0 mm |
| Day 15 | 6 Aug 2022 | Check traps day 2 | 4.6 mm |
| Day 16 | 7 Aug 2022 | Check traps day 3 and apply ZnP bait | 1.2 mm |
| Day 17 | 8 Aug 2022 | | 0.2 mm |
| Day 18 | 9 Aug 2022 | | 0.2 mm |
| Day 19 | 10 Aug 2022 | | 0 mm |
| Day 20 | 11 Aug 2022 | Traps set (Post-baiting population assessment) | 0.4 mm |
| Day 21 | 12 Aug 2022 | Check traps day 1 | 0.2 mm |
| Day 22 | 13 Aug 2022 | Check traps day 2 | 4.2 mm |
| Day 23 | 14 Aug 2022 | Check traps day 3 | 1.4 mm |
| Day 24 | 15 Aug 2022 | Check traps day 4 | 0.4 mm |
| Day 25 | 16 Aug 2022 | Check traps day 5 | 0.4 mm |
| Day 26 | 17 Aug 2022 | Check traps day 6 | 0.2 mm |

*Population sampling to assess mortality*

For pre-baiting population sampling, 20 Longworth traps were placed in each enclosure for three nights (total = 60 trap nights/enclosure; Figure S1). The number of mice per enclosure was reduced but not to less than 10 during the pre-baiting population sampling. This number reflects the moderate mouse densities at which growers are trying to control mice to minimise crop damage and was considered the minimum needed to adequately observe treatment effects and provide statistically robust data.

Trapping effort was increased for the post-baiting population sampling (removal trapping; Figure S1). Trapping ran for six nights (~200 trap nights per enclosure), and on the last two nights, cornflour was dusted over likely mouse burrows to look for active burrows where additional traps were laid. Trapping continued until no mouse activity was observed for two days (as assessed by no captures and undisturbed cornflour on burrows). Captured mice were scanned for PIT tags and measured for weight and head-body length and examined for overall condition and signs of poisoning. Captured mice were examined for overall condition and for signs of poisoning (general alertness: normal to lethargic; movement: normal exploratory activity to abnormal/unbalanced (ataxia) or a reluctance to move; body position: normal to sternal or lateral recumbency; clarity of eyes: bright, clear and open to dull, half or fully closed; respiratory pattern: normal, shallow; gasping; slow, rapid) (Hinds *et al.* 2023; Khan and Schell 2014; Mason and Littin 2003). Mice were then humanely killed by cervical dislocation and then necropsied to assess the condition of major organs (liver, spleen, kidneys, stomach content, lungs) to look for sub-acute signs of toxicity.

*Necropsy of mice found dead*

On necropsy, the eight mice found dead showed signs of mottling and colour change in their livers, consistent with ZnP-affected mouse livers observed in laboratory studies (Hinds *et al.* 2023) and were considered to have died from zinc phosphide poisoning (Table S2).

**Table S2**. Observations of organs after necropsy of mice found dead in enclosures. All were considered to have died from zinc phosphide poisoning. N = normal.

| Enclosure | Sex & ID | Liver | Kidneys | Spleen | Heart | Lungs | Comments |
|---|---|---|---|---|---|---|---|
| Pen 8 (0 kg/ha) | ♀, 200051 | Dark red, mottled | N | N | N | N (bright pink) | |
| Pen 4 (10 kg/ha) | ♀, 200119 | Mottled, pale red/brown | N | N | N | N (bright pink) | |
| Pen 4 (10 kg/ha) | ♂, 362358 | Mottled | N | N | N (but pale) | N (dark pink) | |
| Pen 3 (20 kg/ha) | ♂, 362387 | Mottled, brown/red | N | N | N | N (pink) | Rigor mortis |
| Pen 7 (80 kg/ha) | ♀, 360899 | Red/brown, mottled | N | N | N | N (bright pink) | Rigor mortis |
| Pen 7 (80 kg/ha) | ♀, 360822 | Dark red, some mottle | N | N | Bright red | N (bright pink) | |
| Pen 7 (80 kg/ha) | ♂, 362389 | Dark red, some mottle | N | N | N | N (dark red, uniform) | |
| Pen 9 (320 kg/ha) | ♀, 200050 | N | N | N | N | N (dark pink colour) | |

*Bait toxicity testing*

A sample of 20 grains was randomly taken from the same drum used to bait mice in enclosures and individually analysed to verify bait toxicity by ACS Laboratories [Australia] Pty Ltd (Kensington, Victoria, Australia). Each grain was weighed directly into a Teflon digestion tube to which 5 ml of concentrated nitric acid and 1 ml of 30% hydrogen peroxide were added. The digestion tube was then microwaved for 65 minutes using a temperature gradient from 0-200°C in a Milestone Connect Ethos Lean Compact Microwave Digestor. Once cooled the contents of the digestion tube were diluted to 50 ml in deionised water, followed by a further 10-fold dilution in deionised water for ICP analysis using an Agilent 5110 ICP-OES (Dual View). Data were reported as mg ZnP/grain.
